# TinA enables kinesin-14/KlpA to exhibit processive minus-end-directed motility

**DOI:** 10.1101/2025.02.11.636922

**Authors:** Akasit Visootsat, Andrew R. Popchock, Yuan Gao, Weihong Qiu

## Abstract

Kinesin-14 motors contribute to spindle assembly by localizing to spindle poles and anchoring the minus ends of spindle microtubules. Unlike other kinesin-14 motors, KlpA uniquely exhibits plus-end-directed motility on single microtubules as individual homodimers. However, the mechanism by which KlpA achieves minus-end-directed motility on single microtubules remains elusive. Here, we report that TinA, a highly conserved microtubule-anchoring protein, serves as an activator of KlpA for minus-end-directed motility. TinA directly interacts with KlpA to form minus-end-directed complexes that exhibit continuous movement on microtubules with two distinct velocity modes. The assembly of KlpA-TinA complexes depends on TinA binding to the central stalk of KlpA. Furthermore, TinA is a microtubule-binding protein, with its C-terminal region playing a critical role in microtubule interaction. Deletion of the C-terminus of TinA markedly reduces its microtubule-binding ability and severely impairs the formation of KlpA-TinA complexes. Nonetheless, KlpA-TinA complexes formed without the C-terminus of TinA still exhibit minus-end-directed motility, albeit with a single velocity mode. Collectively, these findings provide critical mechanistic insights into how TinA modulates KlpA, enabling the kinesin-14 motor to achieve minus-end-directed motility.

## INTRODUCTION

The mitotic spindle is a bipolar, microtubule-based structure that eukaryotic cells assemble during mitosis to ensure the accurate segregation of duplicated chromosomes between daughter cells^1^. Microtubules are asymmetric cylindrical polymers of α/β-tubulin heterodimers with two distinct ends: a highly dynamic plus end capped with β-tubulin and a less dynamic minus end capped with α-tubulin^2^. The assembly of a bipolar spindle relies on the coordinated activities of kinesins^3,4^, motor proteins that convert chemical energy from ATP hydrolysis into forces or movements along microtubules^5^. Errors in spindle assembly can result in daughter cells with abnormal chromosome numbers, a hallmark of cancer^6^.

All kinesins contain at least one motor domain that hydrolyzes ATP and binds to microtubules. Based on the sequence similarity of the motor domains, kinesins are classified into 14 subfamilies (kinesin-1 through kinesin-14) and an orphan group^7^. Kinesin-14s are commonly homodimers consisting of an N-terminal microtubule-binding tail, a central coiled-coil stalk, a neck, and two identical C-terminal motor domains^8^. Examples of homodimeric kinesin-14s include HSET from *Homo sapiens*^9^, Ncd from *Drosophila melanogaster*^10,11^, Pkl1 from *Schizosaccharomyces pombe*^12^, and KlpA from *Aspergillus nidulans*^13^. Kinesin-14 motors contribute to spindle assembly by using minus-end-directed motility to localize at spindle poles, anchoring the minus ends of spindle microtubules^8^. Unlike all other kinesin-14 motors studied to date^10,11,14–18^, which either exhibit minus-end-directed motility or diffuse without directional preference, KlpA uniquely displays plus-end-directed motility on single microtubules as individual homodimers^19^. However, it remains unclear how KlpA achieves minus-end-directed motility for localizing to the spindle poles to contribute to spindle assembly. In *S. pombe*, localization of Pkl1 to spindle poles depends on Msd1 and Wdr8^20^, suggesting that KlpA might use a similar mechanism to localize the spindle poles. Notably, Msd1 and Wdr8 are conserved in many eukaryotic organisms^21^, with homologs in *A. nidulans* identified as TinA and AnWdr8, respectively^22^.

In this study, we focused on characterizing the effect of TinA on the motility of KlpA. We found that TinA binds to the central stalk of KlpA, forming KlpA-TinA complexes that exhibit continuous minus-end-directed motility on the microtubules with both fast and slow velocities. We further revealed that the C-terminus of TinA acts as a microtubule-binding tether to not only induce the complexes to move with the slower velocity mode but also play an essential role in the formation of the KlpA-TinA complexes. To our knowledge, TinA is the first protein shown to reverse kinesin directionality through direct interaction. These findings significantly expand our understanding of kinesin regulation by binding partners. Overall, our findings provided critical mechanistic insights into the regulation of KlpA by TinA that enables long-distance minus-end-directed motility on the microtubules.

## RESULTS

### TinA and KlpA interact to form minus-end-directed complexes

We set out to determine whether TinA can directly interact with KlpA to affect its motility on the microtubules. To address this, we first engineered fluorescently labeled KlpA and TinA (GFP-KlpA and TinA-mCherry, Fig. 1a), and then performed an *in vitro* total internal reflection fluorescence (TIRF) microscopy assay to examine the motility of GFP-KlpA on surface-immobilized polarity-marked microtubules, both in the absence and presence of TinA-mCherry (Fig. 1b). In the control motility experiments containing 2.5 nM GFP-KlpA without TinA-mCherry, all but a few GFP-KlpA particles exhibited plus-end-directed motility on the microtubules (Fig. 1c; Supplementary Movie 1), similar to previous observations^19^. The minus-end-directed GFP-KlpA particles (white arrowhead, Fig. 1c) were aggregates rather than individual homodimers, as they were much brighter than the ones moving toward the microtubule plus ends. In contrast, in the motility experiments containing 2.5 nM GFP-KlpA and 80 nM TinA-mCherry, GFP-KlpA molecules completely lost their directional preference for the microtubule plus ends and instead moved continuously toward the minus ends (Fig. 1d, left panel; Supplementary Movie 2). Notably, while TinA-mCherry lacked the ability to autonomously generate directional motility on the microtubules (data not shown), it became motile in the presence of GFP-KlpA and co-translocated with GFP-KlpA toward the microtubule minus ends (Fig. 1d, middle and right panels). Thus, TinA directly interacts with KlpA to form minus-end-directed KlpA-TinA complexes.

**Figure 1:**
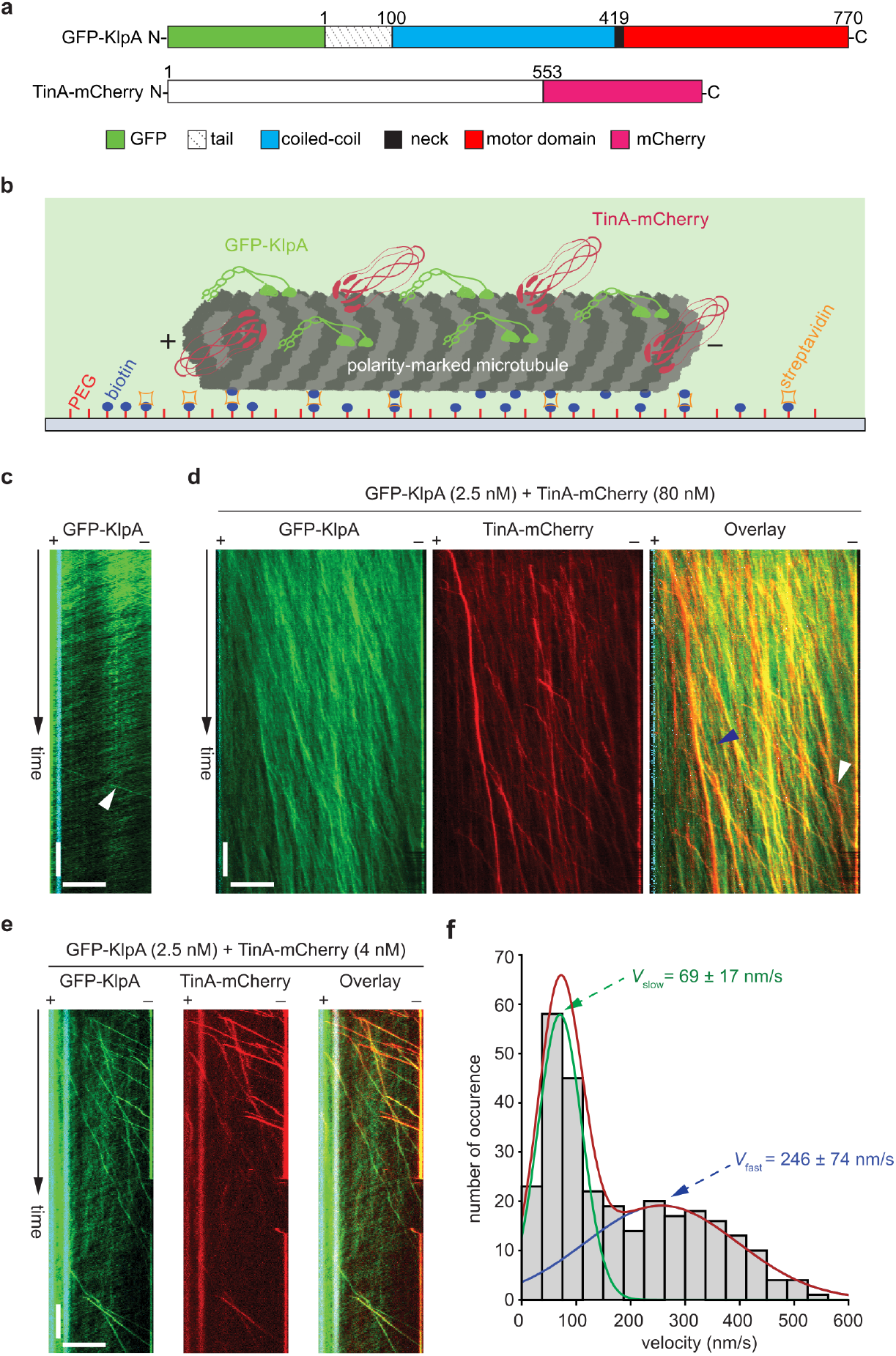
TinA enables KlpA to exhibit minus-end-directed motility with two distinct velocity modes. (**a**) Schematic diagrams of GFP-KlpA and TinA-mCherry. (**b**) The schematic diagram of the in vitro motility assay. (**c**) A representative kymograph showing that in the absence of TinA-mCherry, GFP-KlpA molecules exhibited plus-end-directed motility on a single microtubule. The white arrowhead indicates a GFP-KlpA aggregate moving toward the microtubule minus end. The motility experiment was performed with 2.5 nM GFP-KlpA in the motility chamber. Horizontal scale bar, 10 μm; vertical scale bar, 1 minute. (**d**) Representative kymographs showing that in the presence of a relatively high concentration of TinA-mCherry, GFP-KlpA molecules all exhibited minus-end-directed motility on a single microtubule and co-translocated toward the minus end with TinA-mCherry. The motility experiment was performed with 2.5 nM GFP-KlpA and 80 nM TinA-mCherry in the motility chamber. Horizontal scale bar, 10 μm; vertical scale bar, 1 minute. White arrowheads indicate kymograph segments with fast minus-end-directed motility, and blue arrowhead indicate kymograph segments with slow minus-end-directed motility. (**e**) Representative kymographs showing that in the presence of a much lower concentration of TinA-mCherry, a small number of GFP-KlpA molecules became minus-end-directed and co-translocated with TinA-mCherry toward the minus end on a single microtubule. The motility experiment was performed with 2.5 nM GFP-KlpA and 4 nM TinA-mCherry in the motility chamber. Horizontal scale bar, 10 μm; vertical scale bar, 1 minute. (**f**) The velocity histogram of minus-end-directed KlpA-TinA complexes. The histogram was generated from motility experiments with 2.5 nM GFP-KlpA and 4 nM TinAΔC1-mCherry. The velocity histogram was fitted to a bimodal distribution, as indicated by the red curve. The green and blue curves indicate the slow and fast distributions, respectively.

We observed that minus-end-directed KlpA-TinA complexes appeared to move with two different velocities (Fig. 1d, right panel, blue and white arrowheads). To verify this, we performed additional motility experiments with 2.5 nM GFP-KlpA and 4 nM TinA-mCherry, which allowed us to distinguish individual minus-end-directed complexes and measure their velocities (Fig. 1e; Supplementary Movie 3). Quantitative kymograph analysis confirmed that the velocity distribution of KlpA-TinA complexes was indeed bimodal, with the two modes corresponding to *V*_slow_ = 69 ± 17 nm/s and *V*_fast_ = 246 ± 74 nm/s (mean ± s.d., n = 287, Fig. 1f).

### TinA binds to the central stalk of KlpA

As an initial step toward understanding how the binding of TinA causes KlpA to reverse its direction of motion, we sought to determine where TinA binds on KlpA. We had previously found that KlpA becomes a minus-end-directed motor when its N-terminal nonmotor domain and C-terminal motor domains bind to two different microtubules^19^ or its central stalk is remodeled with the insertion of a flexible polypeptide^23^. These findings led us to hypothesize that TinA binds to the nonmotor region of KlpA. To test this hypothesis, we engineered GFP-KlpAΔN^T^ (Supplementary Fig. 1a) — a GCN4 leucine zipper-driven KlpA tetramer containing an N-terminal GFP and the C-terminal amino acids (a.a.) 398-770 of KlpA — and used TIRF microscopy to examine its motility both in the absence and presence of TinA-mCherry. The results showed that GFP-KlpAΔN^T^ ([GFP-KlpAΔN^T^] = 2.5 nM) exhibited almost identical minus-end-directed motility on the microtubules both in the absence of TinA-mCherry (Supplementary Fig. 1b; Supplementary Movie 4) and in its presence ([TinA-mCherry] = 80 nM; Supplementary Fig. 1c; Supplementary Movie 5). In the latter experiments, no TinA-mCherry was observed to co-translocate with GFP-KlpAΔN^T^ on the microtubules (Supplementary Fig. 1c, middle and right panels). Collectively, these results suggest that the N-terminus of KlpA (a.a. 1-397) is necessary for the interaction between TinA and KlpA.

We next wanted to determine whether the N-terminal region of KlpA (a.a. 1-397) is sufficient for the interaction with TinA. To address this, we engineered GFP-KlpA*Ncd-#1 (Fig. 2a) — a chimera containing an N-terminal GFP followed by the N-terminal a.a. 1-397 of KlpA and the C-terminal motor domains of Ncd (a.a. 305-700) — and examined its motility on the microtubules both in the absence and presence of TinA-mCherry. The motor domains of Ncd were chosen for making GFP-KlpA*Ncd-#1, because GFP-Ncd lacked the ability to interact with TinA-mCherry to form minus-end-directed complexes in the control motility experiments (Supplementary Fig. 1d,e; Supplementary Movie 6, 7). In the motility experiments with 2.5 nM GFP-KlpA*Ncd-#1 in the absence of TinA-mCherry, GFP-KlpA*Ncd-#1 exhibited diffusive motion on the microtubules with no directional preference for either end (Fig. 2b; Supplementary Movie 8). In contrast, in motility experiments with 2.5 nM GFP-KlpA*Ncd-#1 and 80 nM TinA-mCherry, GFP-KlpA*Ncd-#1 gained the ability to generate directional motility on the microtubules and co-translocated with TinA-mCherry continuously toward the minus ends (Fig. 2c; Supplementary Movie 9), similar to what we observed in the motility experiments with 2.5 nM GFP-KlpA and 80 nM TinA-mCherry (Fig. 1d). Thus, the N-terminal region of KlpA (a.a. 1-397) is sufficient for the interaction between TinA and KlpA.

**Figure 2:**
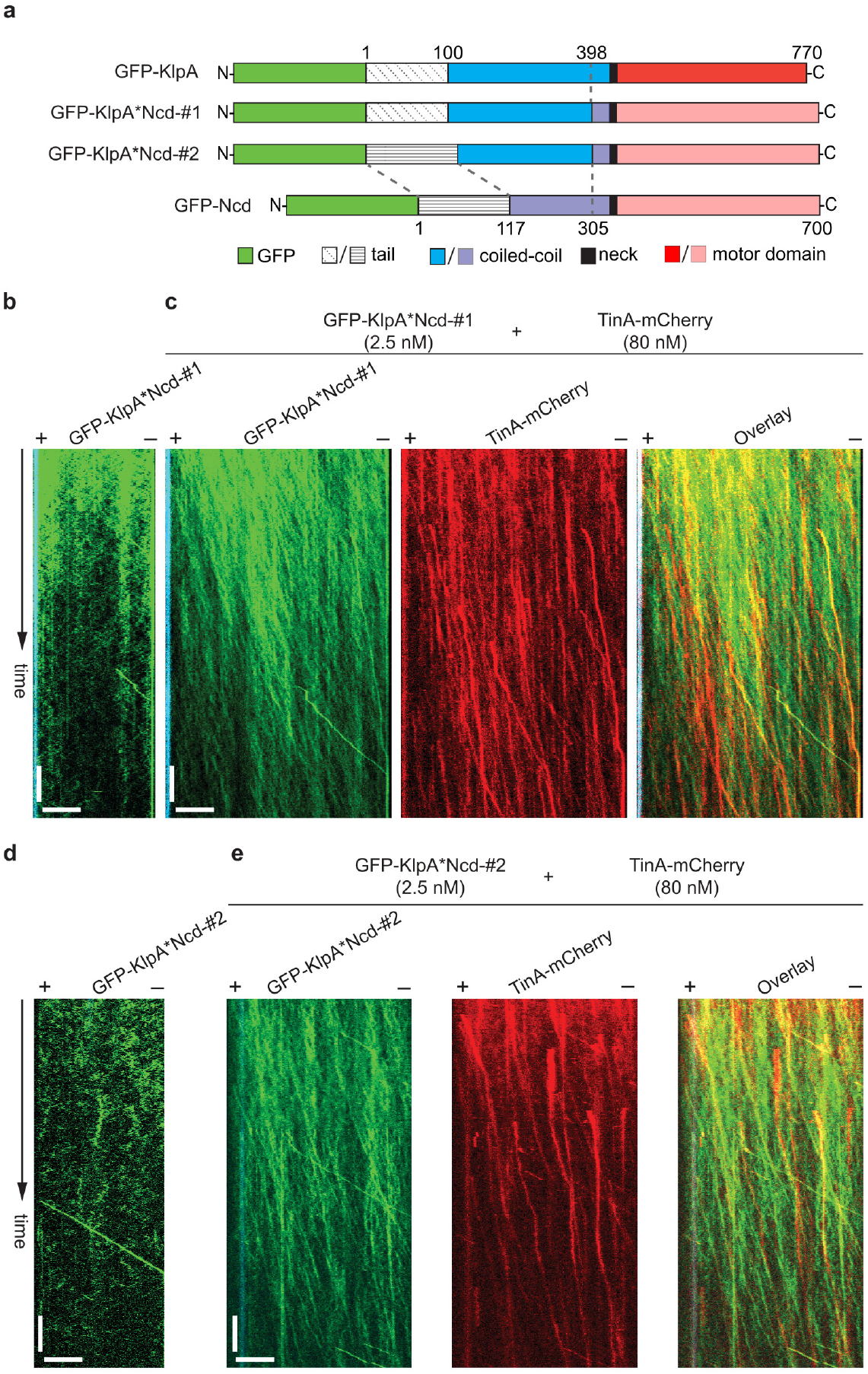
TinA binds to the central stalk of KlpA. (**a**) Schematic diagrams of GFP-KlpA, GFP-KlpA*Ncd-#1, GFP-KlpA*Ncd-#2, and GFP-Ncd. (**b**) A representative kymograph showing that GFP-KlpA*Ncd-#1 exhibited diffusive motion with no directionality preference on a single microtubule. Horizontal scale bar, 10 μm; vertical scale bar, 1 minute. The motility experiment was performed with 2.5 nM GFP-KlpA*Ncd-#1 in the motility chamber. (**c**) Representative kymographs showing that in the presence of a relatively high concentration of TinA-mCherry, GFP-KlpA*Ncd-#1 exhibited minus-end-directed motility and co-translocated with TinA-mCherry. The motility experiment was performed with 2.5 nM GFP-KlpA*Ncd-#1 and 80 nM TinA-mCherry in the motility chamber. Horizontal scale bar, 10 μm; vertical scale bar, 1 minute. (**d**) A representative kymograph showing that GFP-KlpA*Ncd#2 exhibited diffusive motility with no directional preference on a single microtubule. Horizontal scale bar, 10 μm; vertical scale bar, 1 minute. The motility experiment was performed with 2.5 nM GFP-KlpA*Ncd#2 in the motility chamber. (**e**) Representative kymographs showing that in the presence of a relatively high concentration of TinA-mCherry, GFP-KlpA*Ncd#2 exhibited minus-end-directed motility on a single microtubule and co-translocated with TinA-mCherry toward the minus end. The motility experiment was performed with 2.5 nM GFP-KlpA*Ncd#2 and 80 nM TinA-mCherry in the motility chamber. Horizontal scale bar, 10 μm; vertical scale bar, 1 minute.

To further narrow down the region on KlpA involved in the interaction with TinA, we created GFP-KlpA*Ncd-#2 (Fig. 2a), a chimera derived from GFP-KlpA*Ncd-#1 by replacing the region corresponding to the tail of KlpA (a.a. 1-100) with that of Ncd (a.a. 1-117). We found that in the absence of TinA-mCherry, GFP-KlpA*Ncd-#2 ([GFP-KlpA*Ncd-#2] = 2.5 nM) mainly exhibited diffusive motion on the microtubules with no directional preference for either end (Fig. 2d; Supplementary Movie 10). However, in motility experiments with 2.5 nM GFP-KlpA*Ncd-#2 and 80 nM TinA-mCherry, GFP-KlpA*Ncd-#2 and TinA-mCherry were observed to co-translocate continuously on the microtubules toward the minus ends (Fig. 2e; Supplementary Movie 11). Collectively, these results suggest that TinA mainly binds to the central stalk (a.a 100-398) of KlpA to form the minus-end-directed KlpA-TinA complexes.

### The C-terminus of TinA plays a dual-functional role in KlpA-TinA complexes

In the control motility experiments, TinA was observed to bind to microtubules (Supplementary Fig. 1c, middle panel), indicating that TinA functions as a microtubule-binding protein. Microtubule-associated proteins often contain clusters of basic residues within their microtubule-binding regions^24–28^. Consistent with this, we found that the C-terminal region of TinA (a.a. 370–553) contains an abundance of basic residues (Fig. 3a). To validate whether TinA is a microtubule-binding protein and assess the contribution of its C-terminal region, we generated TinAΔC1-mCherry (Fig. 3b), which lacks the C-terminal a.a. 370–553. We then performed in vitro microtubule co-sedimentation experiments with both TinA-mCherry and TinAΔC1-mCherry. Our results showed that TinA-mCherry co-precipitated with microtubules (Fig. 3c, left panel), whereas TinAΔC1-mCherry retained the ability to bind to the microtubules, albeit with a reduced affinity compared to TinA-mCherry (Fig. 3c, right panel). These findings indicate that the C-terminus of TinA constitutes, or at least contributes to, a microtubule-binding motif.

**Figure 3:**
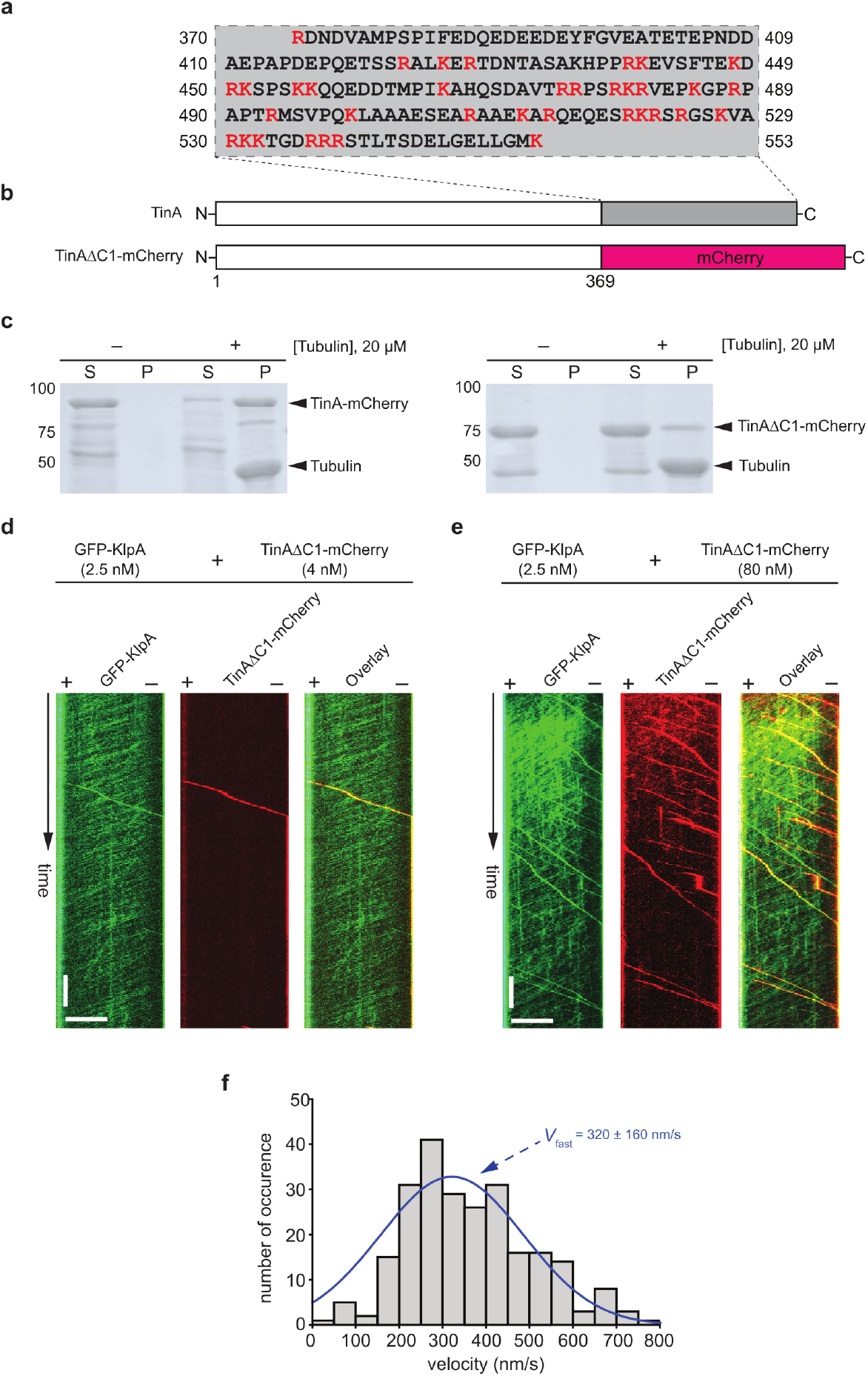
The C-terminus of TinA affects the formation and velocity distribution of KlpA-TinA complexes. (**a**) The C-terminus of TinA (a.a. 370-553) contains an abundance of basic residues. (**b**) Schematic diagrams of TinA and TinAΔC1-mCherry. (**c**) Coomassie-stained SDS-PAGE gels from the microtubule co-sedimentation experiments for TinA-mCherry (Left Panel) and TinAΔC1-mCherry (Right Panel). (**d**) Representative kymographs showing that minus-end-directed KlpA-TinAΔC1 complexes were very rarely observed when the motility experiments were performed with 2.5 nM GFP-KlpA and 4 nM TinAΔC1-mCherry. Horizontal scale bar, 10 μm; vertical scale bar, 1 minute. (**e**) Representative kymographs showing that many more minus-end-directed KlpA-TinAΔC1 complexes were observed when the motility experiments were performed with 2.5 nM GFP-KlpA and 80 nM TinAΔC1-mCherry. Horizontal scale bar, 10 μm; vertical scale bar, 1 minute. (**f**) The velocity histogram of minus-end-directed KlpA-TinAΔC1 complexes. The histogram was generated from motility experiments with 2.5 nM GFP-KlpA and 80 nM TinAΔC1-mCherry. The histogram was fitted to a single Gaussian distribution, as indicated by the blue curve.

To investigate whether the C-terminal region of TinA (a.a. 370–553) influences the motility of KlpA-TinA complexes, we conducted motility experiments with 2.5 nM GFP-KlpA and varying concentrations of TinAΔC1-mCherry. At 4 nM TinAΔC1-mCherry, the motility of GFP-KlpA was largely unaffected, with GFP-KlpA only occasionally observed co-translocating with TinAΔC1-mCherry toward microtubule minus ends (Fig. 3d, Supplementary Movie 12). However, at 80 nM TinAΔC1-mCherry, more TinAΔC1-mCherry molecules were observed to co-translocate with GFP-KlpA toward the microtubule minus end (Fig. 3e and Supplementary Movie 13). Collectively, these motility results suggest that without the C-terminus, the ability of TinA to interact with KlpA to form minus-end-directed complexes is severely impaired but not abolished entirely. Interestingly, KlpA-TinAΔC1 complexes exhibited a single velocity mode comparable to the fast mode of the KlpA-TinA complexes (*V*_*fast*_ = 321 ± 82, mean ± s.d., n = 244, Fig. 3f). This finding implies that the C-terminus of TinA is responsible for the slow velocity mode observed in minus-end-directed KlpA-TinA complexes (Fig. 1d,e).

## DISCUSSION

To summarize, we investigated the effect of TinA on the motility of KlpA, a kinesin14 motor that uniquely exhibits plus-end-directed motility on single microtubules as individual homodimers. We revealed that TinA directly interacts with KlpA, enabling the kinesin-14 motor to reverse its directionality and exhibit continuous motility towards the microtubule minus ends. To our knowledge, TinA is the first protein that has been demonstrated capable of inducing directional reversal in any kinesin through direct interaction. This finding markedly expands current understanding of the scope of kinesin regulation by binding partners.

How does TinA enable KlpA to reverse directionality? We previously showed that KlpA reverses its directionality in two scenarios: either when its N-terminal tail detaches from the microtubule to which its motor domains bind^19^, or when its N-terminal tail and C-terminal motor domains are mechanically decoupled via the insertion of a flexible peptide in the central stalk^23^. Given that TinA binds directly to the central stalk of KlpA (Fig. 2), we suggest that TinA binding alters the structure of the central stalk of KlpA. This alteration may either decouple the N-terminal tail and C-terminal motor domains of KlpA or cause the N-terminal tail of KlpA to detach from the microtubule, thus enabling the kinesin-14 motor to change its directionality. The precise mechanism underlying this process awaits elucidation through future structural studies.

In addition to activating KlpA for minus-end directionality, TinA also enables KlpA to exhibit processive motility on the microtubules (Fig. 1d,e). Our results showed that in the GFP channel, minus-end-directed KlpA-TinA complexes are noticeably brighter than those individual GFP-KlpA homodimers moving in the opposite direction (Fig. 1d), indicating that TinA recruits two or more KlpA molecules to form individual minus-end-directed complexes. Having two or more KlpA molecules is sufficient for these minus-end-directed KlpA-TinA complexes to achieve processive motility on the microtubules, as clustering of as few as two or more nonprocessive kinesin-14 motors has been demonstrated to yield highly processive minus-end-directed motility^29,30^.

We serendipitously discovered that TinA is a microtubule-binding protein (Fig. 3c). The C-terminus of TinA (a.a. 370-553) constitutes, or at least contributes to, a microtubule-binding motif, as removal of the C-terminal region weakens but does not abolish entirely the ability of TinA to bind to the microtubules (Fig. 3b). In principle, TinA can use its C-terminus to enhance the processivity of the resultant KlpA-TinA complexes, as studies have shown that several kinesins gain processive motility via nonmotor microtubule-binding sites^14,17^. Surprisingly, we found that TinAΔC1-mCherry (which lacks the C-terminal a.a. 370-553 and shows reduced microtubule binding) appears to have lost the ability to interact with GFP-KlpA to form minus-end-directed complexes in the motility experiments with 2.5 nM GFP-KlpA and 4 nM TinAΔC1-mCherry (Fig. 3d); but the formation of the minus-end-directed complexes can be partially rescued by increasing the concentration of TinAΔC1-mCherry in the motility experiments (Fig. 3e). Thus, the C-terminus of TinA contributes to enhancing the formation of the KlpA-TinA complexes, but is not involved in the binding interface between TinA and KlpA. How does the C-terminus of TinA contribute to the formation of the KlpA-TinA complexes without directly participating in the binding interface? We suggest that the formation of the KlpA-TinA complexes occurs primarily on the microtubule lattice rather than in the solution (Fig. 4). In this model, the C-terminus of TinA (a.a. 370-553) acts as a primary microtubule-binding tether, which allows TinA to encounter and interact with multiple KlpA molecules on the microtubule lattice to form the KlpA-TinA complexes (Fig. 4a). Without the C-terminal a.a. 370-553, TinAΔC1 is unable to bind to the microtubule as strongly as the wildtype TinA (Fig. 3c, right panel). Consequently, the chance of TinAΔC1 to encounter and interact with multiple KlpA molecules on the microtubule lattice for forming the minus-end-directed complexes is significantly impaired (Fig. 3d,e). Consistent with this model, we found that while KlpA-TinA complexes were readily observed on the microtubules in the motility experiments (Fig. 1d,e), the complexes were not observable in the absence of the microtubules using the standard biochemical methods such as gel filtration and sucrose density gradient centrifugation (data not shown). We further suggest that once KlpA-TinA complexes are formed on the microtubule, the C-terminus of TinA in the complexes can either stay attached to or detached from the microtubule (Fig. 4b). When the C-terminus of TinA in a KlpA-TinA complex is attached to the microtubule (Fig. 4b, top panel), the complex exhibits slow minus-end-directed motility (Fig. 1d); and when the C-terminus of TinA detaches from the microtubule (Fig. 4b, bottom panel), the KlpA-TinA complex exhibits fast minus-end-directed motility (Fig. 3e).

**Figure 4:**
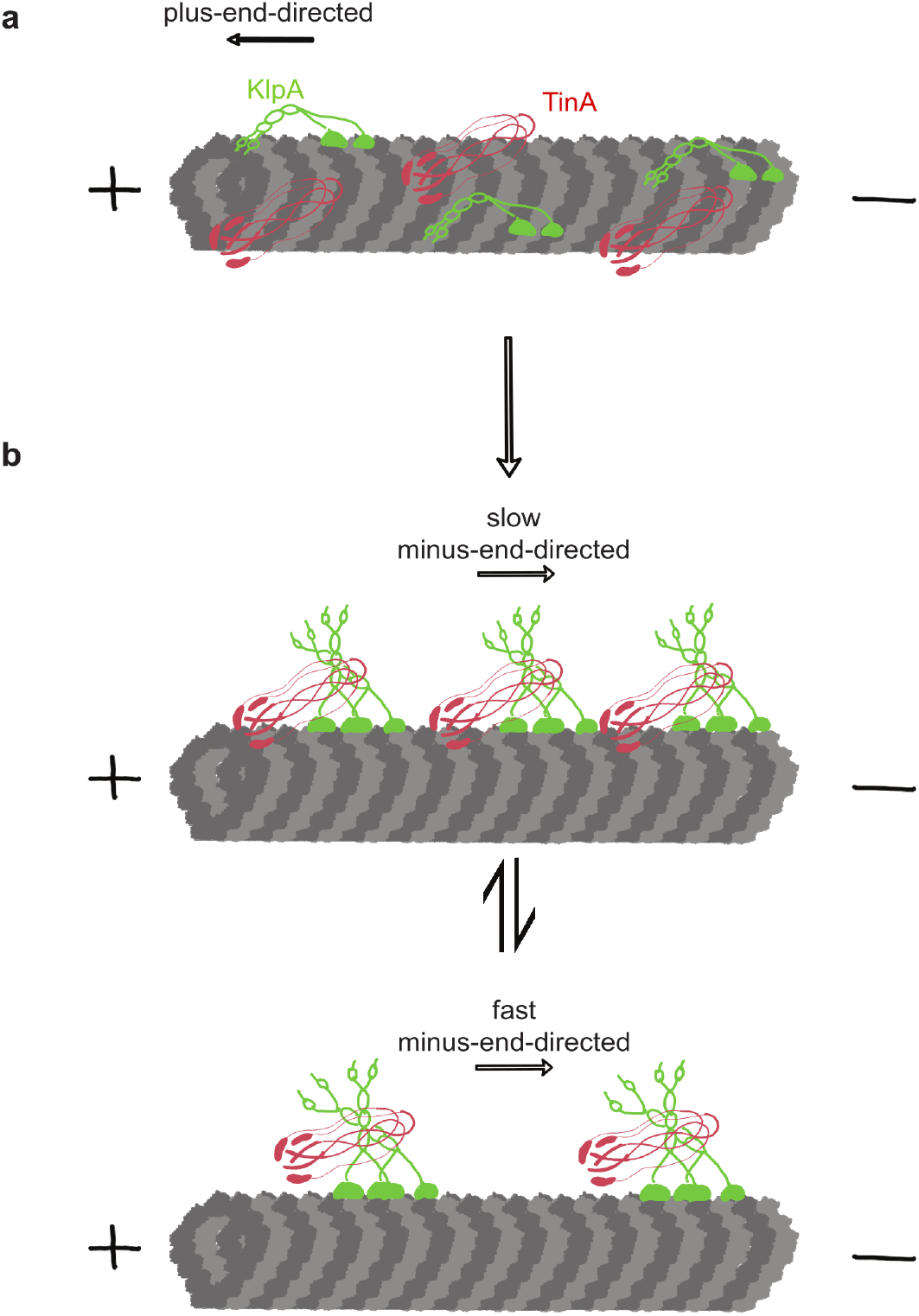
A schematic model for the formation and motility of KlpA-TinA complexes. (**a**) TinA and KlpA interact to form the KlpA-TinA complexes on the microtubule lattice. Prior to interacting with each other to form the KlpA-TinA complexes, KlpA exhibits plus-end-directed motility by binding its N-terminal tail and C-terminal motor domains to the same microtubule, while TinA binds to the microtubule lattice primarily through its C-terminus (a.a. 370-553). (**b**) TinA binds to the central stalk of KlpA to enable the kinesin-14 motor to exhibit minus-end-directed motility with two different velocity modes. The velocity modes of the KlpA-TinA complexes depends on the microtubule-binding state of the C-terminus of TinA. KlpA-TinA complexes exhibit slow minus-end-directed motility when the C-terminus of TinA binds to the microtubule lattice (top panel), but switch to exhibit fast minus-end-directed motility when the C-terminus of TinA detaches from the microtubule lattice (bottom panel).

Our study provides critical mechanistic insights into how TinA enables KlpA to exhibit continuous minus-end-directed motility on microtubules. However, several important questions remain unanswered. One key question pertains to the binding interface of the KlpA-TinA complex. Due to the lack of high-resolution structural data for KlpA, TinA, or the KlpA-TinA complex, we were unable to determine the precise binding interface in this study. Another question involves the role of AnWdr8. In S. pombe, the localization of Pkl1 (a KlpA homolog) to the spindle poles requires both Msd1 (a TinA homolog) and Wdr8^20^. Interestingly, our findings show that TinA alone is sufficient to enable KlpA to exhibit continuous minus-end-directed motility. This observation raises the intriguing possibility that in *A. nidulans*, AnWdr8 may not actively participate in KlpA localization to the spindle poles but instead primarily functions to anchor the kinesin-14 motor at these sites. Additionally, we note that the C-terminus of TinA, while absent in Msd1 from S. pombe, is highly conserved among TinA-like proteins in fungi (Supplementary Fig. 2). This divergence raises important questions about the extent to which the regulatory mechanisms of KlpA and Pkl1 are conserved. Addressing these questions will further enhance our understanding of the regulatory mechanisms governing kinesin-14s during mitotic spindle assembly.

## ACKNOWLEDGEMENTS

This work was supported by the National Institutes of Health (GM127922 to W.Q.).

## AUTHOR CONTRIBUTIONS

W.Q. conceived, designed and supervised the project; A.V. designed and conducted all experiments and performed the analysis; A.R.P. designed and conducted the initial experiments; Y.G. performed the analysis; all authors participated in interpreting the results; and W.Q. wrote the manuscript with the input from other authors.

## METHODS

### Molecular cloning of recombinant KlpA constructs

cDNAs of KlpA, Ncd and TinA were codon-optimized and synthesized (IDT) for enhanced protein expression in bacteria. All recombinant KlpA, Ncd and TinA constructs were integrated into a modified pET-17b vector (Novagen) using either the Gibson assembly cloning kit (NEB) or the Q5 site-directed mutagenesis kit (NEB) and verified by DNA sequencing (Azenta). All protein constructs contained a 6xHis tag (at the N-terminus for all kinesin-14 constructs and at the C-terminus for all TinA constructs) for protein purification. Additionally, all kinesin-14 constructs contained a GFP tag after the N-terminal 6xHis tag, and all TinA constructs contained a mCherry tag before the C-terminal 6xHis tag.

### Protein expression and purification

All protein constructs were expressed in BL21(DE3) Rosetta cells (Novagen). Cells were grown at 37 °C in tryptone phosphate medium (TPM) supplemented with 50 mg/ml Ampicillin until OD600 reached 0.6-0.8. Expression was induced with 0.1 mM isopropyl-b-D-thiogalactoside for 12–14 h at 18 °C. Cells were then harvested by centrifugation, flash-frozen in liquid nitrogen, and stored at −80 °C.

For protein purification, cell pellets were resuspended in 50 mM sodium phosphate (NaPi) buffer (pH 7.2) containing 250 mM NaCl, 1 mM MgCl_2_, 0.5 mM ATP, 10 mM b-mercaptoethanol, 5% glycerol, and 20 mM imidazole in the presence of a protease inhibitor cocktail (phenylmethylsulfonyl fluoride, leupeptin, and pepstatin) and then lysed via sonication. After centrifugation at 27200 *g* for 30 min, soluble protein in the supernatant was purified by Talon resin (Clontech) and eluted into 50 mM NaPi buffer (pH 7.2) containing 250 mM NaCl, 1 mM MgCl_2_, 0.5 mM ATP, 10 mM b-mercaptoethanol, 5% glycerol and 250 mM imidazole. Protein was then flash-frozen in liquid nitrogen and stored at −80 °C. For TinA proteins, ATP was excluded from the purification buffers.

### Preparation of polarity-marked microtubules

Unlabeled tubulin was purified from the porcine brain via the assembly and disassembly method. Polarity-marked microtubules (HiLyte 647) with bright plus ends were prepared as previously described^31^. Briefly, a tubulin mix (containing 1.7 mg/mL unlabeled tubulin and 1.2 mg/mL biotin-labeled tubulin) was first incubated in BRB80 (80 mM PIPES, pH 6.8, 1 mM EGTA, and 1 mM MgCl_2_) with 0.125 mM guanosine-5’-[(a,b)-methyleno]triphosphate (GMPCPP, Jena Bioscience) at 37 °C for 8-10 hours to allow tubulin to polymerize to form microtubules. The resulting tubulin/microtubule mix was then centrifuged using a TLA100 rotor (Beckman) at 279,000 *g* for 7 minutes at 37 °C. To cap the microtubule plus ends, the pellet was resuspended in a bright tubulin mix, which contains 0.75 mg/mL unlabeled tubulin, 0.14 mg/mL HiLyte Fluor647-labeled tubulin (Cytoskeleton) and 1.5 mg/mL N-ethylmaleimide-tubulin in BRB80 with 0.1 mM GMPCPP, and further incubated at 37 °C for additional 45 minutes. The resulting polarity-marked microtubules were first pelleted using a TLA100 rotor at 9800 *g* for 7 minutes at 37 °C and then resuspended in BRB80 with 40 μM Taxol.

### Total internal reflection fluorescence (TIRF) microscopy

All time-lapse imaging experiments were performed at room temperature using the Axio Observer Z1 objective-type TIRF microscope (Zeiss) equipped with a 100x, 1.46 numerical aperture oil-immersion objective and a back-thinned electron multiplier charge-coupled device camera (Photometrics). All experiments used microscope coverslips that were functionalized with biotinylated polyethylene glycol (biotin-PEG) as previously described to reduce nonspecific surface absorption of molecules^32^. All time-lapse imaging experiments in this study used flow cells that were made by attaching a coverslip to a microscope glass slide by double-sided tape.

### In vitro motility assays

For all in vitro motility experiments, the motility chamber was perfused with 0.5 mg/ml streptavidin for immobilizing polarity-marked HyLite 647 microtubules. The chamber was washed twice to remove an unbound streptavidin with buffer A (BRB12 supplemented with 20 μM taxol). The polarity-marked HyLite 647 microtubules were then added and immobilized on the functionalized coverslip for 5 minutes. Unbound microtubules were washed out from the chamber twice with buffer B (BRB50 supplemented with 1.3 mg/ml casein and 20 μM taxol). For the motility experiments of GFP-tagged kinesin-14 molecules (GFP-KlpA, GFP-KlpAΔN^T^, GFP-KlpA*Ncd-#1, GFP-KlpA*Ncd-#2, and GFP-Ncd) in the presence of mCherry-labeled TinA variants (TinA-mCherry and TinAΔC1-mCherry), GFP-tagged kinesin-14 and mCherry-labeled TinA variants were diluted in buffer B and mixed in the motility buffer C (BRB50 supplemented with 25 mM KCl, 1 mM ATP, 25 μM taxol, 1.3 mg/ml casein, and an oxygen scavenger system) for a final concentration of 2.5 nM and either 4 nM or 80 nM, respectively, before added to the chamber. The images were taken immediately after focusing. Time-lapse images were acquired for a total duration of 10 minutes at 1 frame per second with an exposure time of 100 ms for all channels. Kymographs were generated and analyzed using Fiji^33^. The velocity distributions were generated and analyzed in Python^34^.

### Microtubule co-sedimentation assays

For all microtubule co-sedimentation experiments, taxol-stabilized microtubules were prepared as previously described^35^. Briefly, unlabeled-tubulin (20 μM) was incubated on ice for 3 minutes, mixed gently with polymerization buffer (BRB80 supplemented with 2 mM MgCl2, 2 mM GTP, 2 mM DTT, and 20% DMSO), and incubated on ice for additional 3 minutes. The mixture was then incubated at 37 °C for 30 minutes. Polymerized microtubules were mixed with microtubule-stabilizing buffer (BRB80 supplemented with 1 mM DTT and 40 μM taxol) and incubated at 37 °C for additional 30 minutes. Purified TinA-mCherry and TinAΔC1-mCherry were each mixed with taxol-stabilized microtubules in BRB50 supplemented with 25 mM KCl, 1.3 mg/mL casein, 1% ß-mercaptoethanol, and 40 μM taxol. The mixture was incubated at room temperature for 30 minutes and centrifuged at 100000 *g* using a TLA100 rotor for 20 minutes at 25 °C. Coomassie-stained SDS-PAGE gels were used for comparing the amount of protein in the supernatant and pellet fractions.

