## Supplementary Figures for "TinA enables kinesin-14/KlpA to exhibit processive minus-end-directed motility"

### Supplementary Figures 1-2

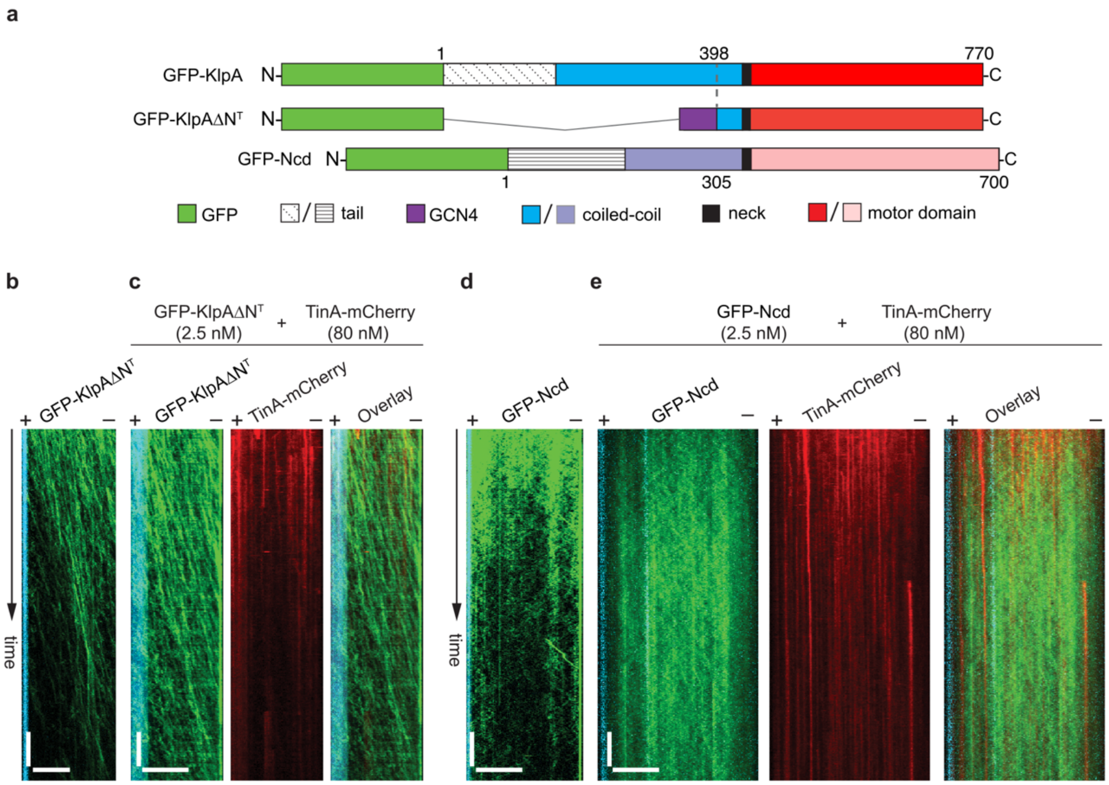


**Supplementary Fig. 1: TinA does not bind to the C-terminal motor of KlpA and the full length Ncd.** (**a**) Schematic diagrams of GFP-KlpA, GFP-KlpAΔN$^T$, and GFP-Ncd. (**b**) A representative kymograph showing that GFP-KlpAΔN$^T$ exhibited minus-end-directed motility in the absence of TinA-mCherry. Horizontal scale bar, 10 μm; vertical scale bar, 1 minute. (**c**) Representative kymographs showing that GFP-KlpAΔN$^T$ retained minus-end-directed motility in the presence of TinA-mCherry and did not co-localize with TinA-mCherry. Horizontal scale bar, 10 μm; vertical scale bar, 1 minute. (**d**) A representative kymograph shows that GFP-Ncd exhibited highly diffusive with no directionality preference in the absence of TinA-mCherry. Horizontal scale bar, 10 μm; vertical scale bar, 1 minute. (**e**) Representative kymographs showing that GFP-Ncd retained its diffusive motility in the presence of TinA-mCherry and did not co-localize with TinA-mCherry. Horizontal scale bar, 10 μm; vertical scale bar, 1 minute.

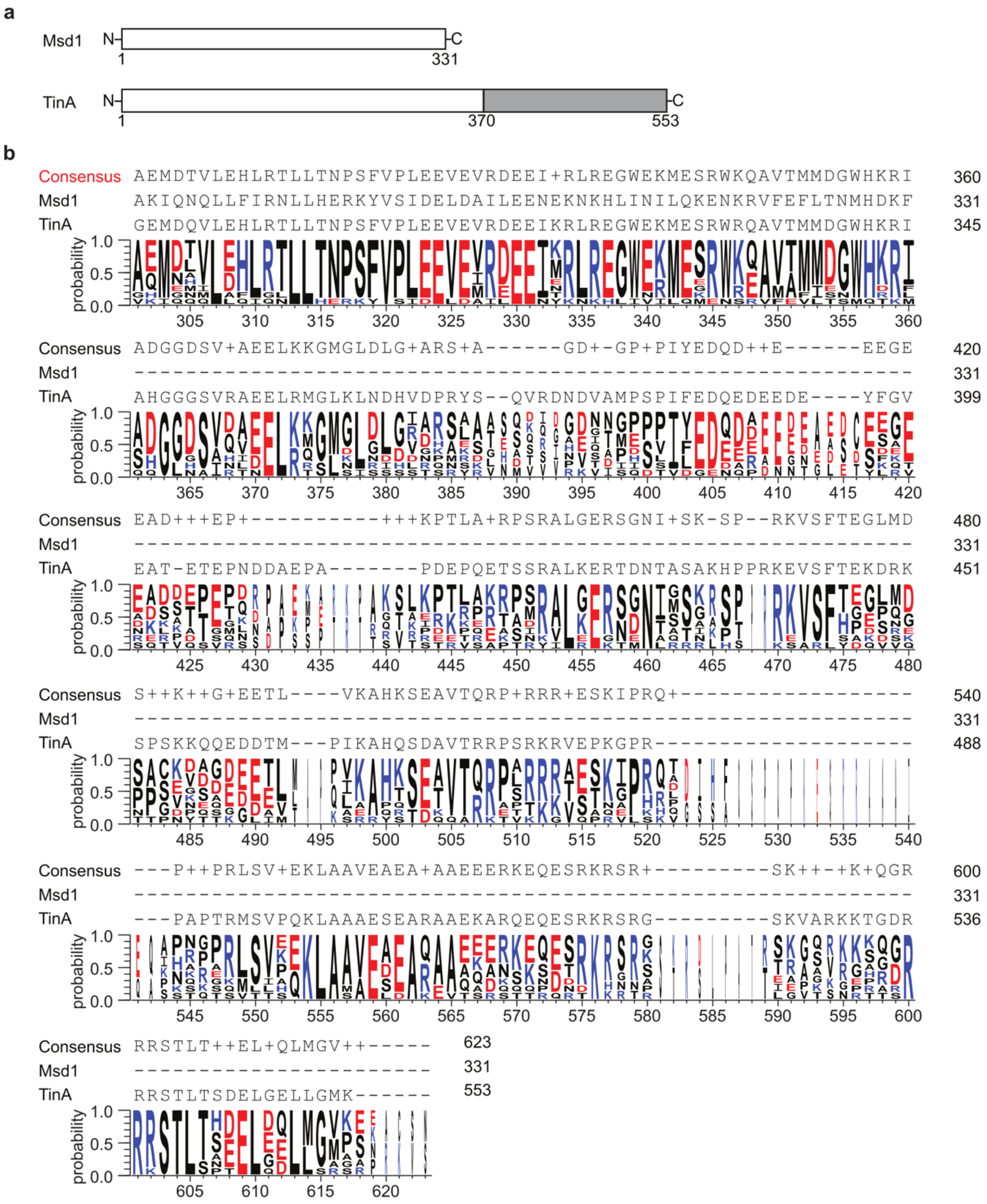


**Supplementary Fig. 2: The C-terminus of TinA is highly conserved among TinA-like proteins in fungi. (a)** Schematic diagrams of Msd1 and TinA showing that TinA contains an extra C-terminal extension. **(b)** The multiple sequence alignment shows that the C-terminal extension of TinA is absent in Msd1 but highly conserved among TinA-like proteins in fungi. The alignment was performed using Clustal Omega in Jalview based on a total of ten input sequences, including Msd1 (UniProt ID: Q9P6R4), TinA (UniProt ID: Q86ZN8), and eight other fungal proteins (Uniprot IDs: A0A100IL75, A0A1V6XGH4, A0A1E3BB44, A0A7H8QNB5, A0A232M1V5, A0A2B7WPF6, A0A0F4YJQ9, and A0A443HMG4).
