## Supplementary Movies for "TinA enables kinesin-14/KlpA to exhibit processive minus-end-directed motility"

**Supplementary Movies 1-13**

**Supplementary Movie 1**

This movie corresponds to Fig. 1c. This movie was acquired with 2.5 nM GFP-KlpA in the absence of TinA-mCherry. Under this condition, GFP-KlpA exhibited plus-end-directed motility and accumulated at the plus end on a single polarity-marked Hilyte 647-microtubule. Top: The microtubule channel, and the arrowhead indicates the microtubule plus end; Middle: The GFP-KlpA channel; Bottom: The overlay of the microtubule and GFP-KlpA channels.

**Supplementary Movie 2**

This movie corresponds to Fig. 1d. This movie was acquired with 2.5 nM GFP-KlpA and 80 nM TinA-mCherry. Under this condition, GFP-KlpA molecules all exhibited minus-end-directed motility on a single polarity-marked Hilyte 647-microtubule and co-translocated with TinA-mCherry toward the microtubule minus end. Top: The microtubule channel, and the arrowhead indicates the microtubule plus end; Second: the GFP-KlpA channel; Third: The TinA-mCherry channel; and Bottom: The overlay of the microtubule, GFP-KlpA and TinA-mCherry channels.

**Supplementary Movie 3**

This movie corresponds to Fig. 1e. The movie was acquired with 2.5 nM GFP-KlpA and 4 nM TinA-mCherry. Under this condition, the majority of GFP-KlpA molecules exhibit the plus-end-directed motility on a single polarity-marked Hilyte 647-microtubule, and individual minus-end-directed KlpA-TinA complexes were observed. Top: The microtubule channel, and the arrowhead indicates the microtubule plus end; Second: The GFP-KlpA channel; Third: The TinA-mCherry channel; and Bottom: the overlay of the microtubule, GFP-KlpA and TinA-mCherry channels.

**Supplementary Movie 4**

This movie corresponds to Supplementary Fig. 1b. This movie was acquired with 2.5 nM GFP-KlpA$\Delta N^{T}$. Under this condition, GFP-KlpA$\Delta N^{T}$ exhibited minus-end-directed motility on a single polarity-marked Hilyte 647-microtubule. Top: The microtubule channel, and the arrowhead indicates the microtubule plus end; Middle: The GFP-KlpAΔN$^{T}$ channel; and Bottom: The overlay of the microtubule and GFP-KlpAΔN$^{T}$ channels.

**Supplementary Movie 5**

This movie corresponds to Supplementary Fig. 1c. This movie was acquired with 2.5 nM GFP-KlpAΔtail$^{T}$ and 80 nM TinA-mCherry. Under this condition, GFP-KlpAΔN$^{T}$ exhibited minus-end-directed motility on a single polarity-marked Hilyte 647-microtubule without interacting with TinA-mCherry. Top: The microtubule channel, and the arrowhead indicates the microtubule plus end; Second: The GFP-KlpAΔN$^{T}$ channel; Third: The TinA-mCherry channel; and Bottom: the overlay of the microtubule, GFP-KlpAΔN$^{T}$ and TinA-mCherry channels.

**Supplementary Movie 6**

This movie corresponds to Supplementary Fig. 1d. This movie was acquired with 2.5 nM GFP-Ncd. Under this condition, GFP-Ncd molecules exhibited diffusive motility with no directional preference on a single polarity-marked Hilyte 647-microtubule. Top: The microtubule channel, and the arrowhead indicates the microtubule plus end; Middle: The GFP-Ncd channel; and Bottom: The overlay of the microtubule and GFP-Ncd channels.

**Supplementary Movie 7**

This movie corresponds to Supplementary Fig. 1e. This movie was acquired with 2.5 nM GFP-Ncd and 80 nM TinA-mCherry. Under this condition, GFP-Ncd exhibited diffusive motility on a single polarity-marked Hilyte 647-microtubule and did not interact with TinA-mCherry. Top: The microtubule channel, and the arrowhead indicates the microtubule plus end; Second: The GFP-Ncd channel; Third: The TinA-mCherry channel; and Bottom: the overlay of the microtubule, GFP-Ncd and TinA-mCherry channels.

**Supplementary Movie 8**

This movie corresponds to Fig. 2b. This movie was acquired with 2.5 nM GFP-KlpA*Ncd-#1. Under this condition, GFP-KlpA*Ncd-#1 exhibited diffusive motility with no directional preference on a single polarity-marked Hilyte 647-microtubule. Top: The microtubule channel, and the arrowhead indicates the microtubule plus end; Middle: The GFP-KlpA*Ncd-#1 channel; and Bottom: The overlay of the microtubule and GFP-KlpA*Ncd-#1 channels.

**Supplementary Movie 9**

This movie corresponds to Fig. 2c. This movie was acquired with 2.5 nM GFP-KlpA*Ncd-#1 and 80 nM TinA-mCherry. Under this condition, GFP-KlpA*Ncd-#1 exhibited minus-end-directed motility on a single polarity-marked Hilyte 647-microtubule and co-translocated with TinA-mCherry toward the minus end. Top: The microtubule channel, and the arrowhead indicates the microtubule plus end; Second: The GFP-KlpA*Ncd-#1 channel; Third: The TinA-mCherry channel; and Bottom: the overlay of the microtubule, GFP-KlpA*Ncd-#1 and TinA-mCherry channels.

**Supplementary Movie 10**

This movie corresponds to Fig. 2d. This movie was acquired with 2.5 nM GFP-KlpA*Ncd-#2. Under this condition, GFP-KlpA*Ncd-#2 molecules exhibited diffusive motility with no directional preference on a single polarity-marked Hilyte 647-microtubule. Top: The microtubule channel, and the arrowhead indicates the microtubule plus end; Middle: The GFP-KlpA*Ncd-#2 channel; and Bottom: The overlay of the microtubule and GFP-KlpA*Ncd-#2 channels.

**Supplementary Movie 11**

This movie corresponds to Fig. 2e. This movie was acquired with 2.5 nM GFP-KlpA*Ncd-#2 and 80 nM TinA-mCherry. Under this condition, GFP-KlpA*Ncd-#2 molecules exhibited minus-end-directed motility on a single polarity-marked Hilyte 647-microtubule and co-translocated with TinA-mCherry toward the minus end. Top: The microtubule channel, and the arrowhead indicates the microtubule plus end; Second: The GFP-KlpA*Ncd-#2 channel; Third: The TinA-mCherry channel; and Bottom: the overlay of the microtubule, GFP-KlpA*Ncd-#2 and TinA-mCherry channels.

**Supplementary Movie 12**

This movie corresponds to Fig. 3d. This movie was acquired with 2.5 nM GFP-KlpA in the presence of 4 nM TinAΔC1-mCherry. Under this condition, very few minus-end-directed KlpA-TinAΔC1 complexes were observed. Top: The microtubule channel, and the arrowhead indicates the microtubule plus end; Second: The GFP-KlpA channel; Third: The TinAΔC1-mCherry channel; and Bottom: the overlay of the microtubule, GFP-KlpA and TinAΔC1-mCherry channels.

**Supplementary Movie 13**

This movie corresponds to Fig. 3e. This movie was acquired with 2.5 nM GFP-KlpA and 80 nM TinAΔC1-mCherry. Under this condition, more KlpA-TinAΔC1 complexes were observed on a polarity-marked Hilyte 647-microtubule. Top: The microtubule channel, and the arrowhead indicates the microtubule plus end; Second: The GFP-KlpA channel; Third: The TinAΔC1-mCherry channel; and Bottom: The overlay of the microtubule, GFP-KlpA and TinAΔC1-mCherry channels.
